# European Robin Cryptochrome-4a Associates with Lipid Bilayers in an Ordered Manner, Fulfilling a Molecular-Level Condition for Magnetoreception

**DOI:** 10.1101/2024.07.20.604329

**Authors:** Marta Majewska, Maja Hanić, Rabea Bartölke, Jessica Schmidt, Henrik Mouritsen, Karl-Wilhelm Koch, Ilia A. Solov’yov, Izabella Brand

**Author notes:** Corresponding author; Ilia A. Solov’yov. Izabella Brand **Email:**. These authors contributed equally (Marta Majewska, Maja Hanić, Rabea Bartölke).

## Abstract

Since the middle of the 20th century, long-distance avian migration has been known to rely partly on the geomagnetic field. However, the underlying sensory mechanism is still not fully understood. Cryptochrome 4a (ErCry4a), found in European Robin (*Erithacus rubecula*), a night-migratory songbird has been suggested to be a magnetic sensory molecule. It is sensitive to external magnetic fields via the so-called radical-pair mechanism. ErCry4a is primarily located in the outer segments of the double cone photoreceptor cells in the eye, which contain stacked and highly ordered membranes that could facilitate the anisotropic attachment of ErCry4a needed for magnetic compass sensing. Here, we investigate possible interactions of ErCry4a with a model membrane that mimics the lipid composition of outer segments of vertebrate photoreceptor cells by using experimental and computational approaches. Experimental results show that the attachment of ErCry4a to the membrane could be controlled by the physical state of lipid molecules (average area per lipid) in the outer leaflet of the lipid bilayer. Furthermore, polarization modulation infrared reflection absorption spectroscopy allowed us to determine the conformation, motional freedom, and average orientation of the α- helices in ErCry4a in a membrane-associated state. Atomistic molecular dynamics studies supported the experimental results. A ∼1000 kcal mol^−1^ decrease in the interaction energy as a result of ErCry4a membrane binding was determined compared to cases where no protein binding to the membrane occurred. At the molecular level, the binding seems to involve negatively charged carboxylate groups of the phosphoserine lipids and the C-terminal residues of ErCry4a. Our study reveals a potential direct interaction of ErCry4a with the lipid membrane and discusses how this binding could be an essential step for ErCry4a to propagate a magnetic signal further and thus fulfill a role as a magnetoreceptor.

## Introduction

Every year, millions of birds migrate and navigate precisely across the globe, covering long distances between their breeding and wintering grounds.^1^ In the middle of the 19^th^ century, it was suggested that migratory birds might use the geomagnetic field for navigation^2^, but it took until 1966 before Merkel^3^ could demonstrate it scientifically. Over the years, it has been revealed that migratory birds have an intrinsic magnetic inclination compass that requires light of specific wavelengths to operate properly.^4^ The magnetic compass was found to be present in the eyes of the birds^4–6^, and magnetic compass information is processed in a specific part of their visual system.^7–10^ Furthermore, electromagnetic radiation of a broad range of frequencies disrupts the birds’ ability to orient using their magnetic compass.^11–18^

One self-consistent explanation for magnetic field sensing by migratory birds is provided by a radical pair mechanism that can occur inside proteins.^7, 19–25^Radicals are transient states of molecules with an odd number of electrons. Under certain conditions, two such radicals can form spin correlated pairs, where the unpaired electron spins appear to be either parallel or anti- parallel, forming triplet or singlet states, respectively.^19, 20, 23, 26–31^ The radical pair mechanism hypothesis suggests that non-equilibrium dynamics change the population of singlet and triplet states inside the bird’s eye and that this process is affected by the external geomagnetic field.^20, 21^ In the context of the avian magnetoreception, the radical-pair mechanism was first proposed by Schulten and colleagues^32^, who furthermore, in the year 2000, suggested cryptochromes (Cry) as a candidate protein to host the radical pairs needed for the operation of the magnetic compass inside a cell.^22^

Most birds have three different cryptochrome genes, which give rise to at least six different isoforms (Cry1a, Cry1b, Cry2a, Cry2b, Cry4a, Cry4b).^6, 33–42^ Some cryptochromes bind a flavin-adenine dinucleotide (FAD) cofactor which is required for the formation of radical pairs inside the protein.^20, 42, 43^ In birds, it seems that Cry1 and Cry2 do not bind FAD, whereas purified recombinant ErCry4a binds FAD and is sensitive to external magnetic fields *in vitro*. ^42, 44, 45^ Furthermore, Cry1 and Cry2 are extremely conserved across all phylogenetic groups of birds ^46^, whereas Cry4a is characterized by strong positive selection and high variability which are typical characteristics of sensor proteins.^46^ In the bird retina, ErCry4a is specifically expressed in the outer segments of double-cone and long-wavelength single-cone photoreceptor cells^5^, and ErCry4a expression increases during the migratory season.^5^ All these characteristics make ErCry4a the most promising primary magnetic receptor candidate.^5, 20, 42^ Pigeon ClCry4 shows similar photochemistry and radical pair formation,^37, 47^ but it seems to differ from ErCry4a with respect to localization in the retina and protein-protein interactions.^37^ Correspondingly, it is not clear whether pigeons can use a light-dependent radical-pair-based magnetic compass at all as, for instance, neuronal activation of Cluster N has only been seen in night-migratory songbirds, but never in day-active birds.^8, 9, 48^

For efficient sensing of the direction of the Earth’s magnetic field, a magnetoreceptor molecule (e.g. cryptochrome with its radical pair) must be anchored at the cellular and molecular level and ideally show some degree of alignment.^21, 31^ ErCry4a is a cytoplasmic protein with no predicted transmembrane regions. It is expressed in the outer segments of double cone photoreceptor cells, which contains a stack of parallel membrane units called lamellae and thus could provide the orientational order and regularity required for efficient magnetoreception.^5, 21, 35^ The membrane lamellae are separated by ∼15-20 nm and could indeed anchor and align ErCry4a. Therefore they are the focal point for searching of binding partners of ErCry4a.^35, 49, 50, 42^ A yeast-two-hybrid screening identified six potential binding partners of ErCry4a, including iodopsin, a transmembrane protein specific for double cones and long-wavelength single cones, and a specific transmembrane potassium channel, strengthening further the hypothesis of the membrane association of the cryptochrome protein.^49^ The cytosolic α-subunit of the double cone-specific heterotrimeric G-protein was later confrimed as an ErCry4a binding partner, and it seems to be myristoylated at its N-terminus.^49, 51^ Thus, the G protein α-subunit could also act as a membrane anchor and/or as a part of a signaling cascade.^51^ Other ways to anchor a cytoplasmic protein to membranes are acylation or alkylation, electrostatic interactions, protein binding to specific polar residues, hydrophobic interactions, or anchorage of a protein via an amino acid fragment, thereby facilitating the association with a lipid membrane.^52–55^

Common phospholipids such as phosphatidylcholine (PC), phosphatidylethanolamine (PE), phosphatidylserine (PS), sphingomyelin, and cholesterol (chol) are present in the outer segment membranes of vertebrate photoreceptor cells. ^56–58^ Specific for outer segment membranes is the presence of phospholipids with unsaturated acyl chains. Except in the chicken^49^, the exact lipid composition of the outer segment of double cone photoreceptor cells in birds is unknown. The lipid composition maintains the membrane lamellae in a liquid-disordered state. In the liquid state, the average area per phospholipid molecule is usually controlled by the packing of the acyl chains, allowing for water accumulation between the polar head groups. Such molecular arrangement affects the elasticity modulus, surface energy, mechanical stress, and/or membrane curvature.^53, 59^ These physical properties of the membranes, in turn, control important biological functions such as signal transduction, transport of matter, or binding of proteins (e.g., enzymes). ^53, 60–62^

In the present study, we fabricated a single bilayer model of the vertebrate outer segment membrane to investigate whether lipids could contribute to the association of ErCry4a to membranes. Protein-lipid binding was further studied by combining experimental and theoretical approaches. A lipid mixture composed of PC, PE, and PS, all with dimyristoyl chains and chol, was used to construct a fluid lipid membrane (see Section S1 in the supporting information (SI)). In our experiments, one leaflet of the lipid membrane was exposed to the interaction with ErCry4a expressed in E. coli ^35, 42^ and dissolved in an aqueous solution and finally transferred as a free-standing, floating lipid bilayer onto a gold surface (see Fig. S2). Photon-based spectroscopy methods, such as infrared spectroscopy, are particularly attractive to gain information about the conformation and structure of biomolecules present in aqueous solution near the membrane surface.^63–65^ To detect weak IR absorption signals from a single lipid bilayer and an associated protein, the reflection-based technique, polarization modulation infrared reflection absorption spectroscopy (PM IRRAS), was used. The surface selection rule^66^ of IRRAS allowed us to determine the average order parameter (orientation) of ErCry4a in the membrane-associated state. Additionally, we reproduced the experimental conditions computationally and studied the likely interactions between the model lipid membrane and the ErCry4a protein. The simulations were done for two different protein orientations, with either the C- or the N-terminus facing the membrane surface. The results reveal that ErCry4a can associate with a model lipid membrane reaching a uniform, partially restricted orientation as required for sensing the Earth’s magnetic field.^31^

## Materials and Methods

### ErCry4a Protein Expression and Purification

Wild type ErCry4a (GenBank: KX890129.1) was expressed and purified according to the protocol described in detail previously, ^42^ with the following modifications: His-tagged ErCry4a was expressed in BL21(DE3) *Escherichia coli* cells in the dark using lysogeny broth media containing 10 g l^−1^ yeast extract instead of 5 g l^−1^, and the expression time was extended from 22 h to 44 h. Protein samples were purified under red light conditions by Ni-NTA agarose columns, followed by anion exchange chromatography as described previously. Purified protein samples were concentrated to 5-6 mg ml^−1^ in an aqueous buffer solution (20 mM Tris, 250 mM NaCl, 20% glycerol, and 10 mM 2- mercaptoethanol) and 0.6 M Trehalose (Merck KGaA, Darmstadt, Germany) when samples were lyophilized. Samples were either used fresh or snap frozen in liquid nitrogen, lyophilized, and stored at −20°C until the measurements for a maximum of four weeks. For the measurements, the protein sample was buffer exchanged into the electrolyte solution (25 mM *d*11-Tris, 100 mM NaCl, with or without 5 mM MgCl2 in D2O, pD = 7.35) using ZebaTM spin desalting columns, 7K MWCO (Thermo Fisher, Waltham, MA, USA).

### Chemicals

The following lipids were used to fabricate the model lipid membranes: 1,2-dimyristoyl-*sn*-glycero-3-phosphocholine, DMPC (14:0-14:0), 1,2-dimyristoyl-*sn*- glycero-3-phosphoethanolamine, DMPE (14:0-14:0), 1,2-dimyristoyl-*sn*-glycero-3-phospho-L-serine, DMPS (14:0-14:0) (all from Avanti Polar Lipids, USA), and cholesterol from Sigma-Aldrich (Germany). D2O was purchased from Deutero (Germany). Tris(hydroxymethyl)amino- methane, Tris Pufferan® ≥99.9%, Roth (Germany), per-deuterated tris-(hydroxymethyl-d3)- amino-d2-methane *d*11-Tris, 1-thio-*β*-D-glucose, sodium deuteroxide, NaOD, deuterium chloride, DCl, MgCl2 (anhydrous, ≥98%), H2SO4 (puriss. p.a), CHCl3 were purchased from Sigma-Aldrich (Germany), NaCl (≥99.5%) from Roth (Germany), HCl, 25% from AnalaR Normapur, VWR (France), hydrogen peroxide (30%) from Fisher Chemicals (Belgium). Ethanol and methanol were purchased from AnalaR Normapur, VWR (France). All aqueous solutions were prepared using freshly filtered water with conductivity ≤ 0.50 µS cm^-1^ (PureLab Classic, Elga LabWater, Germany).

### Preparation of the floating lipid membrane

A Au(111) disc (15 mm diameter MaTecK, Germany) was left for self-assembly for 20 h in 2 mM 1-thio-*β*-D-glucose in water. After the self-assembly, the thioglucose-modified Au(111) was rinsed with a large amount of water and dried in a stream of argon. Then, a lipid bilayer was deposited on the 1-thio-*β*-D-glucose monolayer modified Au(111) surface using Langmuir-Blodgett and Langmuir-Schaefer transfer, see Fig. S2. The phospholipids were dissolved in the following solvents: DMPC and cholesterol in CHCl3, DMPE in CHCl3:MeOH 7.5:1 *v*/*v*, and DMPS in CHCl3:MeOH:H2O 39:8:1 *v*/*v*. Stock solutions containing fixed mole fractions of each lipid: DMPC (χ = 0.4), DMPE (χ = 0.4), DMPS (χ = 0.15), and cholesterol (χ = 0.05) were mixed to obtain the final 0.65 mg mL^−1^ (1 mM) lipid solution. This lipid mixture mimics the lipid composition of the double-cone outer membrane of vertebrae.^58, 67, 68^ The lipid solution was stored at 4 °C for no longer than 2 weeks. The lipid solution was placed at the liquid|air interface of the Langmuir KSV LB mini trough (KSV, Finland) using a glass microsyringe (Hamilton, USA). The electrolyte solution contained either 25 mM Tris, 100 mM NaCl, 5 mM MgCl2, pH = 7.65 ± 0.05, or 25 mM Tris and 100 mM NaCl, pH = 7.65 ± 0.05. The surface pressure (*Π*) vs. area per molecule (*A*m) isotherms were recorded in a Langmuir trough equipped with two hydrophilic barriers. The accuracy of measurements was ± 0.02 nm^2^ for *A*m and ± 0.1 mN m^−1^ for *Π*. The inner leaflet of the model membrane was transferred onto *β*-Tg modified Au(111) surface by a vertical withdrawing of the substrate through the air|liquid interface at *Π* = 30 mN m^−1^ at a speed of 40 mm min^−1^, see Fig. S2A. The transfer ratio was 1.10 ± 0.10. After the transfer, the monolayer was left overnight for drying. For the transfer of the outer leaflet, the lipid monolayer was compressed to either *Π* = 10 mN m^−1^ or *Π* = 20 mN m^−1,^ and ErCry4a was injected underneath the monolayer to yield 100 nM of the protein in the electrolyte solution. The lipid monolayer was left for 30 min to allow for interaction. The temperature of the aqueous subphase during the interaction of the protein with the lipid monolayer was controlled and was set to either 21°C or 24 °C. Afterward, the lipid-ErCry4 monolayer was compressed at the barrier speed of 10 mm min^−1^ to 25 mN m^−1^, see Fig S2 B. The second outer leaflet of the membrane was transferred using LS transfer, yielding a floating bilayer mimicking the structure of the outer segment double cone photoreceptor cells, see Figure S2 C. Model membranes prepared in this procedure make no contact to the solid substrate and “float” on the thioglucose surface ensuring that the lipids have physiological mobility, being an ideal model to study lipid-protein interactions, see Fig. S2C.

### Polarization modulation infrared reflection absorption spectroscopy

PM IRRA spectra were recorded using a Vertex 70 spectrometer with a photo-elastic modulator (*f* = 50 kHz; PMA 50, Bruker, Germany) and demodulator (Hinds Instruments, USA). A home-made thin electrolyte layer spectroelectrochemical cell for PM IRRAS experiments was used. In this setup, an incoming IR beam passes through an optical window, a thin layer of a liquid, analyzed film, and is reflected from a metal (mirror) surface. The glass spectroelectrochemical cell was rinsed with water, ethanol, and water and dried in an oven at 60 °C overnight. A CaF2 optical window was rinsed with ethanol, water, and ethanol, then dried under an argon stream and placed in a UV ozone chamber (Bioforce Nanosciences, USA) for 10 min. A Au(111) electrode (diameter 15 mm) modified by the floating model membrane was used as the working electrode and mirror for the IR radiation. A platinum counter electrode was built into the spectroelectrochemical cell. The reference electrode was Ag|AgCl in 3 M KCl in D2O. The electrolyte solution was 25 mM *d*11-Tris, 100 mM NaCl, with or without 5 mM MgCl2, pD = 7.35 (the difference between pD and pH is approximately 0.4) in D2O.^69^ The electrolyte solution was deaerated for 30 min by purging with argon. At each potential applied to the Au(111) electrode, 400 spectra at a resolution of 4 cm^−1^ were collected. For the amide I vibrational mode analysis, the half-wave retardation was set to 1600 cm^−1,^ and the angle of incidence of the IR radiation was 64°. The thickness of the electrolyte layer between the prism and the working electrode ranged between 6 and 8 µm. During the measurement, potentials were cycled between 0.1 and −0.4 V. The potential step was −0.1 V in the negative and 0.1 V in the positive-going potential scans. The PM IRRA spectra were analyzed using the OPUS v5.5 software (Bruker, Germany). Two sets of experiments on the model membrane prepared at different temperatures and in the presence or absence of MgCl2 were done.

### Attenuated total reflection infrared spectroscopy (ATR IRS)

64 scans of ATR background and sample spectra were collected with a resolution of 4 cm^−1^. First, the spectra were measured from a drop (15 µl) of the electrolyte solution (25 mM *d*11-Tris, 100 mM NaCl, 5 mM MgCl2 in D2O) (background spectrum) placed on a diamond prism (MVP-Pro ATR unit, Harrick Scientific Products, Inc., USA) using a Bruker Vertex 70 spectrometer. Next, the ATR spectrum was recorded from a drop (15 µl) of the analyte, 4.8 µM ErCry4a, in the electrolyte solution. The subtraction of the analyte spectrum from the background spectrum gave the ErCry4a spectrum.

### Computational methods

An atomistic multi-component system consisting of the ErCry4a protein and the model membrane was constructed using VMD^70, 71^, and CHARMM-GUI^72–74^. ErCry4a protein and the model membrane were first simulated separately, as listed in Table 1, and later merged into one system and simulated jointly. Full-length ErCry4a protein was modeled using AlphaFold^75^ and simulated in three NPT stages and an NVT stage to ensure an equilibrated structure. The atomistic structure of ErCry4a contains 8,591 atoms. Additionally, 0.1 M NaCl was added into the system to be consistent with the experiments. The size of the simulation box containing the ErCry4a protein was 128 Å × 90 Å × 87 Å. The simulations were performed using NAMD^70, 76, 77^, which were interfaced through the VIKING platform.^78^ CHARMM36 force field with CMAP corrections was employed for all simulations.^79–81^ The investigated ErCry4a was set to be in the inactive state, usually present in dark conditions, characterized by the FAD cofactor being in a fully oxidized state. FAD cofactor parameters were adopted from earlier studies.^42, 43, 82–86^

**Table 1.**
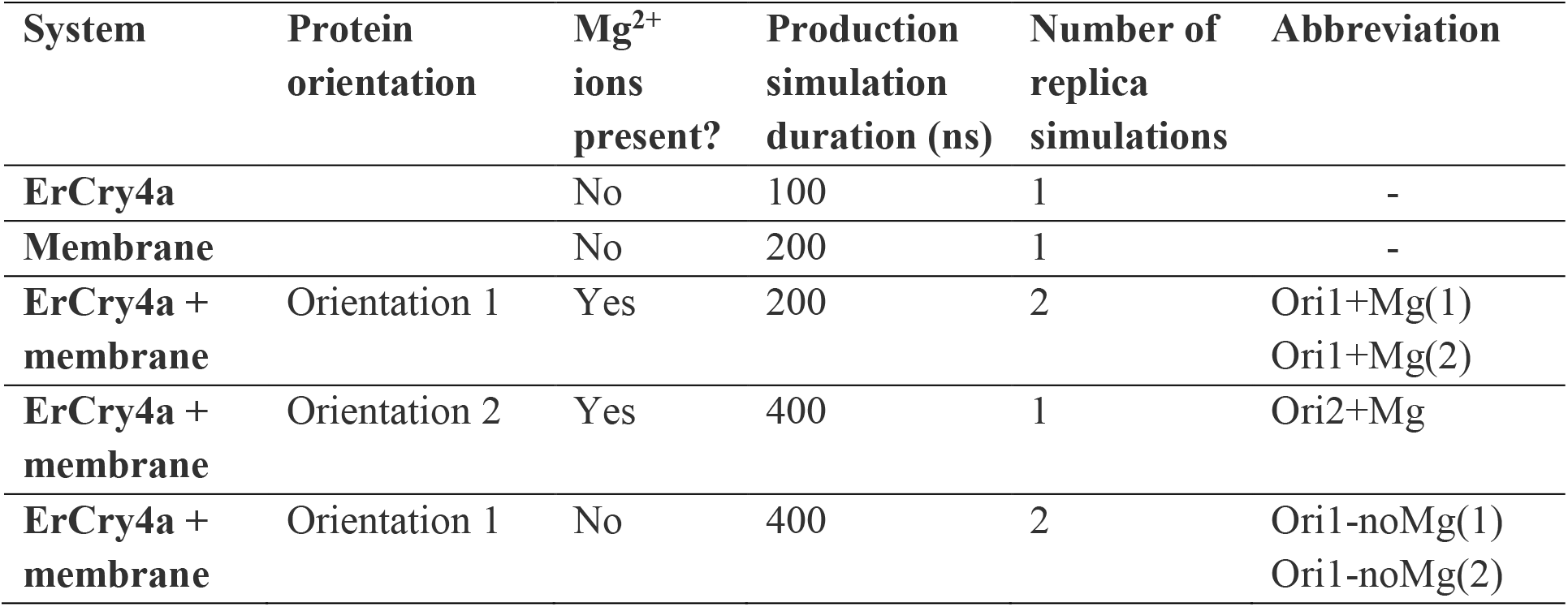
Summary of the performed simulations. Explicit water was present in all simulations.

The membrane was constructed using the CHARMM-GUI Membrane Builder^72–74^ according to the experimental setup consisting of 620 lipid molecules in both monolayers. Molecules that make up the model membrane include DMPC (χ = 0.4), DMPE (χ = 0.4), DMPS (χ = 0.15) and cholesterol (χ = 0.05), see section S1 in the SI. The number of atoms in the membrane bilayer system solvated in a water box was 240,017, where 170,961 atoms correspond to the water molecules. The simulation box size with the membrane was 128.6 Å × 128.6 Å × 142.3 Å at the end of the 200 ns equilibration phase. The membrane thickness and area per lipid were used to estimate the quality of the computationally constructed membrane.

An equilibrated membrane and an equilibrated ErCry4a protein were put together in a simulation box of size 128.6 Å × 128.6 Å × 198.9 Å, and different ErCry4a orientations were probed. In all considered cases, the water molecules in the system were relaxed in a short 1 ns NPT simulation before the production simulations. Two initial protein orientations were considered to test the influence of different orientations of the protein with respect to the membrane. They are referred to as Ori1 and Ori2 in the following text, see Table 1. Ori1 features the C-terminal part of the ErCry4a protein to be closer to the membrane, while Ori2 has the N-terminus of the protein closer to the membrane. To test the influence of Mg^2+^ ions on ErCry4a binding, an additional simulation setup included Ori1 without the presence of the Mg^2+^ ions. These 3 scenarios were simulated for 200 – 400 ns, as summarized in Table 1.

Interaction energies between the protein and the membrane were calculated with the NAMD-energy module of VMD.^71^ The particle-mesh Ewald (PME) summation method was used to treat long-range electrostatic interactions.^87^ Edge-to-edge distances were analyzed via an in-house script. Secondary structure analysis was performed with the STRIDE algorithm^88^ using VMD^89^. To determine the types of secondary structures, STRIDE considers both hydrogen bonding patterns and the backbone geometry of the protein.

### Specific lipid-protein interaction analysis

The PyLipID program was utilized to determine the types of lipids that show interactions with the ErCry4a protein.^90^ PyLipID is a Python package for identifying and characterizing specific lipid-protein interactions. Using output data from MD simulations, it employs the so-called dual cut-off scheme that enables differentiation between lipid conformational rearrangements and dissociation of the protein from the membrane. Additionally, the hydrogen bonding analysis was performed via the HBonds Plugin implemented in VMD.^70, 77^

## Results

ErCry4a needs to be able to accumulate at interfaces in order to associate with a lipid membrane. When dissolved in an aqueous solution, the purified recombinant ErCry4a forms a monolayer at the air|water interface that reaches an equilibrium spreading pressure of 13 mN m^-1^, see Fig S3. The ability of the protein to accumulate at interfaces suggests that ErCry4a may associate, in an ordered manner, at the surface of a biological membrane, being a necessary condition for efficient sensing of the Earth’s magnetic field.^21, 31^

Lipids are amphiphilic molecules and, when deposited at the air|water interface, form stable, ordered monolayers with their polar head groups oriented into the water and the acyl chains oriented towards the air, see schemes in Fig.1 and Fig. S2. Langmuir isotherms provide information about the physical state and packing of the lipids in monolayer films^91^ by depicting the changes of the surface pressure versus area available per molecule. A monolayer of lipids at the air-water interface is the simplest model of one leaflet of a biological membrane. Cell plasma membranes have a surface pressure of ca. 25 – 30 mN m^−1^.^92^ In a monolayer (bilayer) assembly, the presence of lipids with mono- and polyunsaturated acyl chains reduces the surface pressure of the lipid films as compared to molecules with saturated chains of the same length.^91^ The outer segments of photoreceptor cells contain lipids with unsaturated acyl chains, which do not allow for tight packing of the lipids in the membrane; they grant the fluidity and plasticity required for deformations of the membrane forming cone’s lamellae and reduce the membrane surface pressure.^58, 93^ Lipids with unsaturated chains are unstable when exposed to air. Therefore, using lipids with shorter (e.g., C12, lauroyl or C14, myristoyl) saturated chains offers an excellent alternative to form a liquid model membrane of the outer segments of cone photoreceptor cells. Figure 1A shows the surface pressure (*Π)* vs area per molecule (*A*m) isotherms of the DMPC:DMPE:DMPS:chol (0.4:0.4:0.15:0.05 mole ratio) lipid monolayer at the air-electrolyte interface recorded at 21 and 24 °C. The two isotherms have a similar shape and display a temperature dependent change in the slope of the *Π* vs *A*m curves at a mean area per molecule of ca. 0.6 nm^2^. It corresponds to a phase transition from a liquid-disordered (Ld) at low surface pressures to a liquid-ordered (Lo) physical state of the acyl chains at high surface pressures. In the liquid-ordered state, the acyl chains adopt a fully stretched, all-trans conformation.^91, 94^ In the liquid-disordered state the acyl chains “melt” adopting gauche conformations which take more space and prevent a tight packing of the lipid molecules allowing for the accumulation of water in the polar head group region. At the temperature of *T* = 21 °C, the lipid monolayer exists in a liquid disordered state at *Π* < 13 mN m^−1^, while at *T* = 24 °C, it happens at *Π* < 22 mN m^−1^. The calculated compressibility modulus further confirms the existence of the two physical states of the monolayer, see Fig. S4.

**Figure 1.**
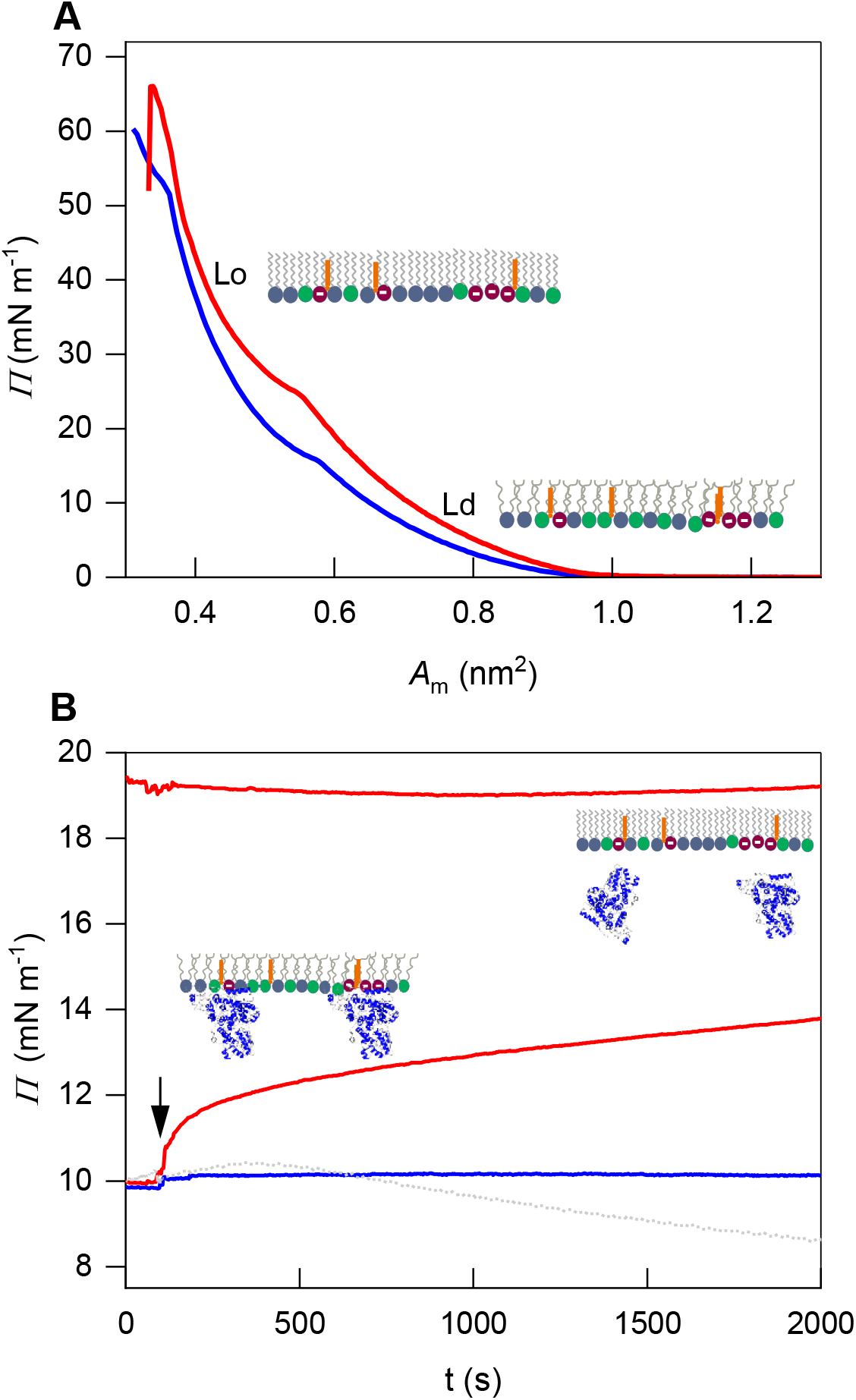
**A**: Surface pressure (*Π)* vs area per molecule (*A*m) isotherms of the lipids in the model membrane of outer segments at the air-electrolyte interface recorded at 21 °C (blue) and 24 °C (red). The insets illustrate the chain order in liquid disordered (Ld) and liquid ordered (Lo) states. **B**: Time dependence of the surface pressure recorded for the lipid monolayers after injection of 100 nM ErCry4a (arrow); Prior to the ErCry4a injection, the lipid monolayer was compressed to the surface pressure of either 10 or 20 mN m^−1^; Line color: 21 °C (blue) and 24 °C (red). Electrolyte solution contained 25 mM *d*11-Tris, 100 mM NaCl, and 5 m MgCl2 in H2O. Grey line: control experiment: changes in the surface pressure of a pure lipid monolayer, in the absence of the protein. Insets illustrate the chain order in the monolayer on the protein attachment.

To test the ErCry4a binding to a lipid film, lipid monolayers at the air-electrolyte interface were compressed to the Ld (*Π* = 10 mN m^−1^) and Lo (*Π* = 20 mN m^−1^) states, respectively. After injection of the ErCry4a into the electrolyte solution phase under the lipid monolayer, the changes in the surface pressure over time were recorded, see Fig. 1B. At 21 °C, after injection of ErCry4a under the monolayer compressed to *Π* = 10 mN m^−1^, the *Π* value did not noticeably change over time (see Fig 1B, blue line). Once the monolayer was compressed to *Π* = 10 mN m^−1^ at 24 °C, an increase in the *Π* value to ∼14 mN m^−1^ occurred after ca. 30 min interaction (see Fig. 1B, red line), indicating membrane association of ErCry4a. Indeed, the attachment of a protein to the membrane usually causes an increase in the surface pressure.^95^ At 24 °C, the increase in the *Π* value did not affect the liquid-disordered state of the monolayer, since the pressure was below the lipid phase transition pressure of 22 mN m^−1^. At 21°C, in contrast, a 3-4 mN m^−1^ increase in *Π* would cause a phase transition of the lipid monolayer to a liquid ordered state, see Fig. S4, inhibiting ErCry4a association to the membrane. No membrane association of ErCry4a to a lipid monolayer existing in a liquid-ordered state was observed at 24 °C once the lipid monolayer was initially compressed to 20 mN m^−1^ (see Fig 1B, red line). The results demonstrate that a liquid-disordered state of the lipid monolayer is crucial for ErCry4a binding, indicating that the lipids’ physical state and thermotropic behavior govern the lipid-ErCry4a interactions. These results demonstrate that the subtle changes in the lipid order and membrane fluidity may drastically alter biological processes on the membrane surface.^53, 60^

The experimental results described above indicate that ErCry4a associates with a lipid monolayer. A model of a membrane of the lamellae in the outer segment of the double cone photoreceptor cell with attached protein was transferred onto a gold surface to study the conformation and alignment of ErCry4a. A floating membrane supported on a thioglucose monolayer self-assembled on a gold surface was transferred by the LB-LS method, see Fig. S2. A floating membrane is an excellent model to investigate the protein–membrane interactions because the lipid molecules do not have any contact with a solid surface and retain their mobility and physical state comparable to those found in a monolayer at the air-water interface. The gold surface efficiently reflects the IR radiation^64^, allowing for the collection of PM IRRA spectra from the floating membrane, which permits the analysis of the protein conformation and average orientation. The amide I vibrational mode of the protein, includes predominantly the C=O stretching mode in the amide group and carries information about the hydrogen bond network at the carbonyl group providing information about the secondary structure elements (α-helices, β-sheets and other structural elements) of proteins.^63^ The amide I mode has to be deconvoluted to gain information about the protein structure. The deconvolution analysis was done for a solution ATR IR spectrum of ErCry4a in D2O, see Fig. 2A and for membrane-associated ErCry4a, see Fig 2 B-D.

**Figure 2.**
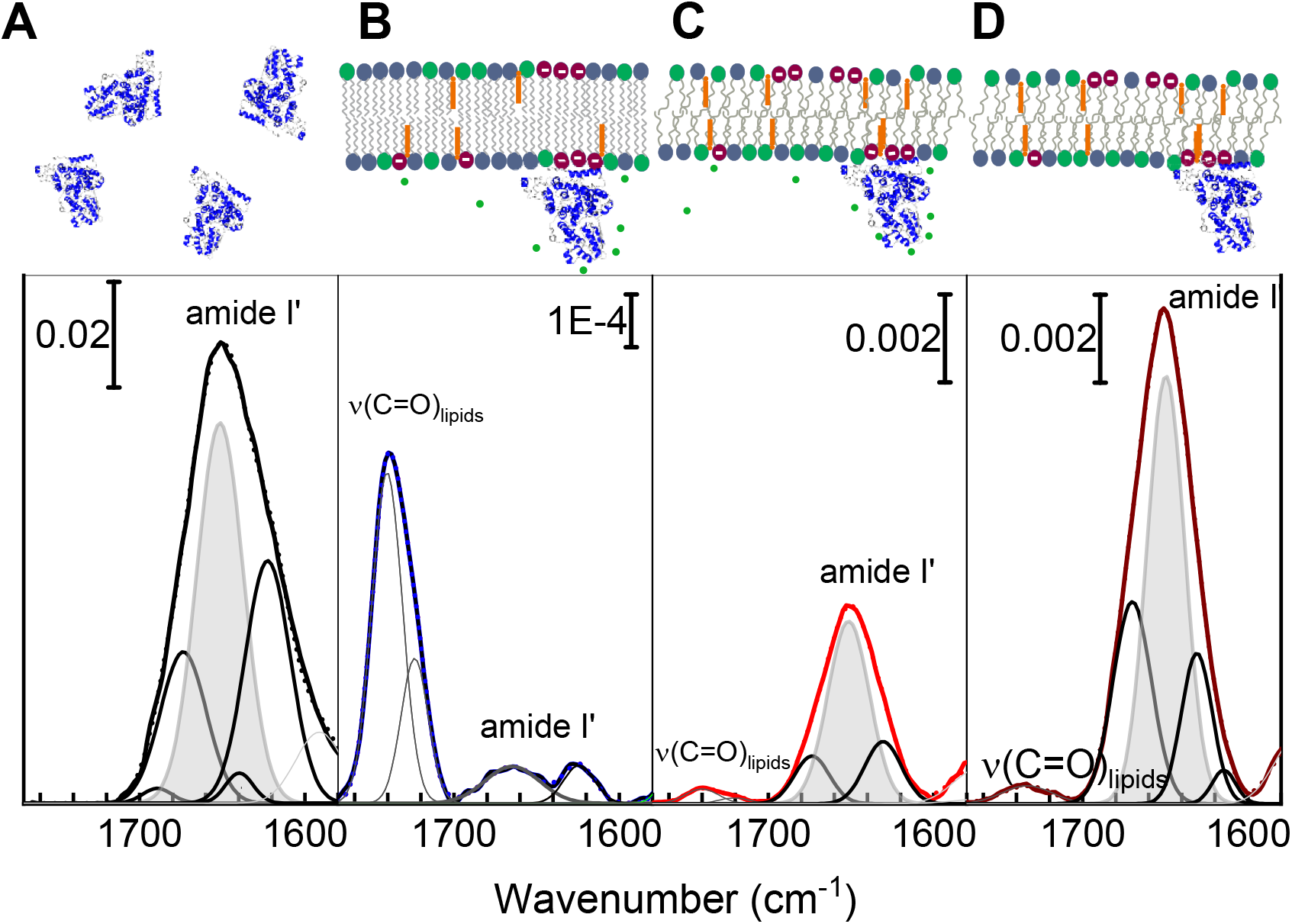
**A**: Deconvoluted ATR IR spectrum (black thick line) of 4.8 µM ErCry4a in D2O; **B-D**: PM IRRA spectra of the model ErCry4a/bilayer system deposited on the Au surface in at **B**: 21 °C (blue thick line), **C**: 24 °C (red thick curve); Electrolyte solution contained 25 mM d11- Tris, 10 mM NaCl and 5 mM MgCl2 in D2O and **D**: at 24 °C (dark red thick line); Electrolyte solution contained 25 mM *d*11-Tris, 100 mM NaCl in D2O. Spectra are shown in the 1780-1580 cm^-1^ region. Thin black and gray lines show the results of deconvolution while the shaded area corresponds to the amide I’ band of the α-helices. Insets illustrate the experimental conditions: Scale bars correspond to absorbance values measured in arbitrary units.

Figure 2A shows a deconvoluted ATR IR spectrum of ErCry4a in D2O. Inserting a protein in D2O causes a slight shift of the amide I mode compared to H2O.^63^ Because the OH deformation band in water overlaps with the amide I band, most of the IR studies of proteins are done in D2O; in this case, the nomenclature changes to amide I’ mode. The second derivative and Fourier self-deconvolution methods were used to deconvolute the amide I’ absorption band. The deconvolution results revealed that the amide I’ band in ErCry4a contains five components centered at 1689 cm^−1^, 1675 cm^−1^, 1653 cm^−1^, 1639 cm^−1^, and 1623 cm^−1^, see Table S1. For ErCry4a in D2O, the most intense and analytically important amide I’ band is centered at 1653 cm^−1^ and attributed to the α-helices, see the shaded area in Fig. 2A. The α- helices constitute 50 ± 2 % of the secondary structure elements in ErCry4a.

Figure 2 shows that the association of ErCry4a to the lipid bilayer affected the intensity and the shape of the amide I’ mode, indicating membrane-association-dependent alternations in the protein folding, conformation, and/or anisotropic orientation. For clarity of interpretation, the ErCry4a amide I’ mode and the ϖ(C=O) of the phospholipids in the model membrane are shown together in Fig. 2 B-D. Independent of temperature and electrolyte composition, the intensity of the ϖ(C=O) mode in the lipids is constant. On the other hand, temperature significantly impacts the amide I’ mode of ErCry4a. At 21 °C, the intensity of the amide I’ mode was weaker than the intensity of the ϖ(C=O) mode in the lipid bilayer, indicating that only a tiny amount of the ErCry4a molecules were bound to the membrane surface, see Fig. 2B. In this case, the amide I’ mode was a single, broad band centered at 1666 cm^−1^, see Table S1. The position of this band is not characteristic for any of the secondary structure elements in proteins^63^, indicating that ErCry4a associating with a liquid-ordered membrane at 21 °C lost its native structure (fold). At 24 °C, independently of the presence of Mg^2+^ ions in the electrolyte solution, an intense amide I’ band centered at 1650 cm^−1^ was detected in the PM IRRA spectra, see Fig. 2 C, D, and Table S1. Note that in the absence of Mg^2+^, an additional amide I’ band at 1614 cm^−1^ was detected, indicating the presence of aggregated β-sheets and pointing at protein aggregation (Fig. 2D).

The positions of the deconvoluted amide I’ modes in the PM IRRA spectra recorded for ErCry4a associated with the lipid bilayers existing in a liquid-disordered state (24 °C) were comparable to those found in the spectrum obtained for the protein in solution, see Table S1. The protein predominantly comprises α-helices and contains some β-sheets and other structural elements (turns and coils). However, the PM IRRAS experiments show an enhancement of the signal recorded from the α-helices, see Fig. 2C, D and attenuation of signals from other structural elements. The experimentally observed enhancement of the amide I’ mode attributed to the α-helices does not indicate protein denaturation but occurs due to an anisotropic orientation of the protein upon association with the membrane. According to the surface selection rule of IRRAS ^66^, the intensity of an IR absorption mode in an anisotropic film depends on the surface concentration of the adsorbed species and on the orientation of the transition dipole moments in the electric field of the reflected IR radiation, see Section S7 in the SI. The latter condition leads to the enhancement of the amide I’ mode from the α-helices in ErCry4a associated with the membrane.

PM IRRAS experiments were done under electrochemical control at different potentials applied to the gold electrode, thus at different membrane potentials. Recent findings indicate that a light-induced deformation of the membrane’s lamellae in the outer segments of photoreceptor cells changes the structure and composition of the electrical double layer at the membrane surface, affecting the membrane potential.^96, 97^ We have therefore tested if the electric potentials affect membrane-associated ErCry4a. No potential-dependent spectral changes were detected in the amide I’ mode, indicating that the electric potentials and, thus, the membrane potentials do not affect the conformation and orientation of membrane-associated ErCry4a.

A uniformly ordered protein film attached to a lipid membrane is expected to reduce the motional freedom of the proteins significantly. The motional order of a protein fragment, e.g. α-helices, represents the number of degrees of motional freedom expressed by a particular moiety or molecular fragments and is typically described through the order parameter (*S*).^98^ Referencing the integral intensities of the deconvoluted amide I’ bands of ErCry4a bound to the membrane to the intensities of the solution spectrum of the protein allows the quantitative analysis of the average orientation of different structural elements (e.g. α-helices) in ErCry4a associated with the lipid membrane. A previously described procedure^99, 100^ was used to calculate the average order parameter (*S*0) attributed to the long axes of all α-helices in the membrane-associated proteins.

The average angle ‹*θ*› between the transition dipole moments of the amide I’ vibrational modes arising in all 26 helices in ErCry4a and the surface normal (direction of the electric field of the reflected IR beam) is defined as

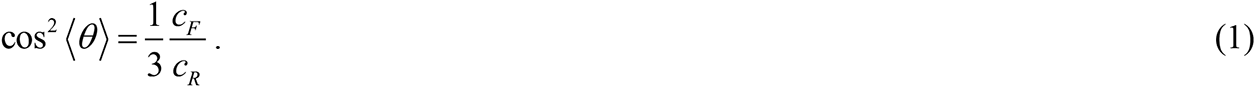

corresponds to the percent content of α-helices in the membrane associated ErCry4a.

This value was obtained from the ratio between the integral intensity of the amide I’ band of the α-helix band and the integral intensity of the entire amide I’ band. *c_R_* = 50 ± 2% and corresponds to the content of α-helices in the solvated ErCry4a (random distribution). The dependency of *θ* on the applied electric potential is shown in Fig. S5. The angle *θ* could further be used to calculate the order parameter *S*helix of the of the amide groups in ErCry4a helices as (see Fig. S6):

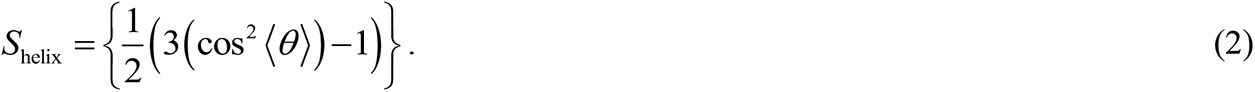

The order parameter *S*0 for the long axes of the α-helical fragments in ErCry4a is then defined as:

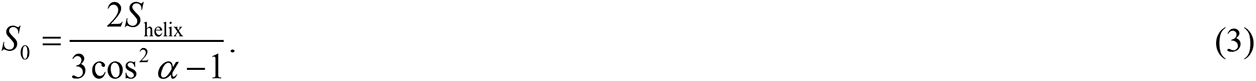

Here, α is the characteristic angle between the long axis of an α-helix and the transition dipole moment of the amide I’ vibration mode. This angle is known to be 34 °– 38 °; see Section S7 and Fig. S7.^101–103^ In the solution phase, the protein is expected to have a random orientation resulting in *S0* = 0. The experimentally determined *S*0 for membrane-associated ErCry4a is shown in Fig. 3.

**Figure 3.**
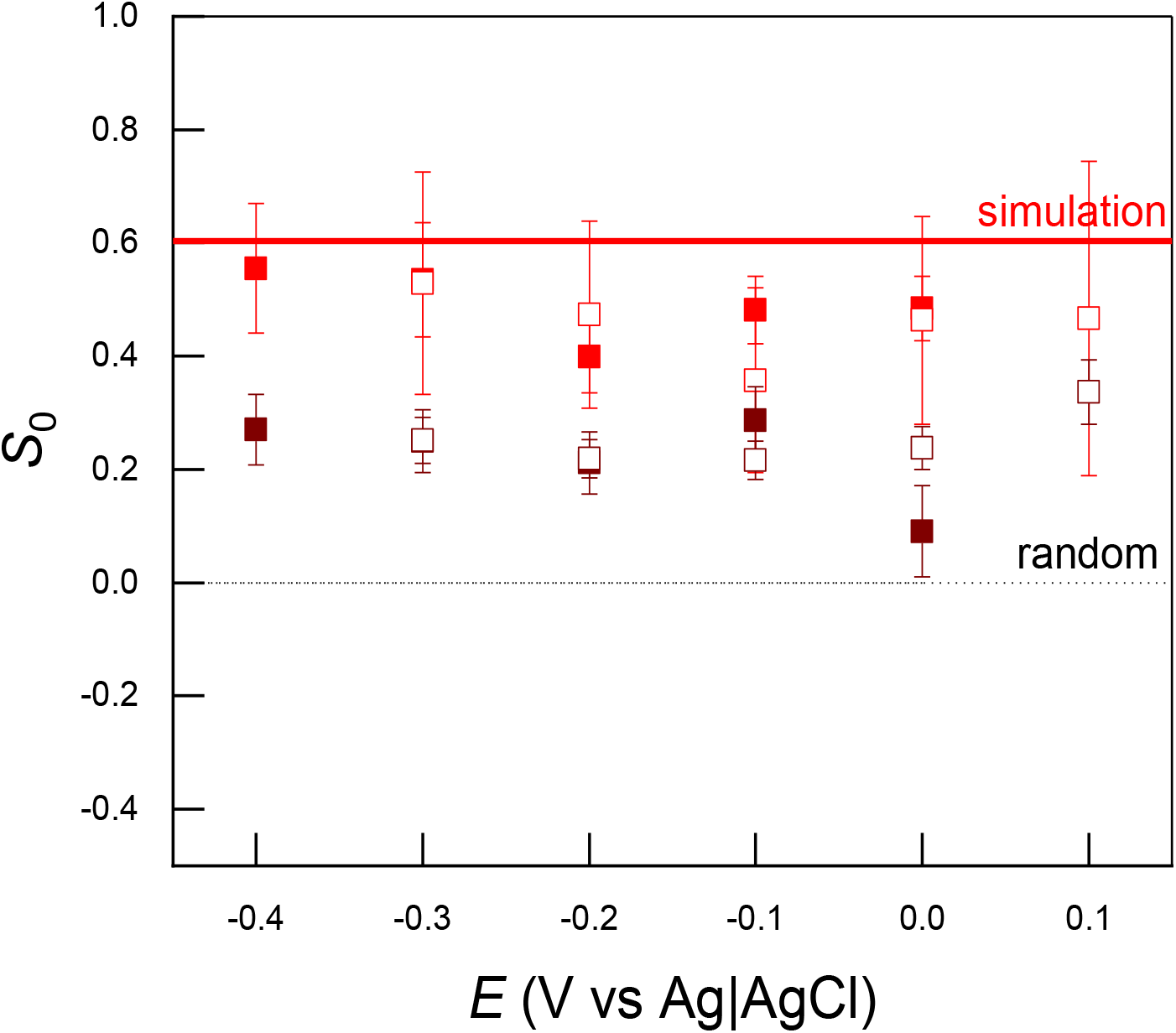
Order parameter (*S0*) of the long axis of all α-helical fragments in ErCry4a measured at different values of the electrode potential for the ErCry4a protein associated with the lipid bilayer deposited on the Au surface at 24 °C in the presence of 5 mM MgCl2 (red symbols) and absence of Mg^2+^ (dark red) in the 25 mM *d*11-Tris, 100 mM NaCl in D2O electrolyte solution; opened and closed symbols correspond to the positive and negative going potential scans. Solid red line shows the order parameter determined from the simulations, while the dashed line indicates the order parameter for randomly oriented proteins.

At 24 °C, the quantitative analysis reveals differences in the average orientation of the α- helices in the membrane-associated ErCry4a in the presence and absence of Mg^2+^. Independently of the electric potential, in the presence of Mg^2+^, *S*0 is lower and equals 0.47 ± 0.07, while in the absence of Mg^2+^, it was lower and equaled 0.23 ± 0.07. The *S*0 value calculated from the MD simulations resulted in *S*0 = 0.60 ± 0.05 and was computed for a single ErCry4a protein associated with the lipid membrane (see text below). Both experimental and computational results confirmed a uniform ordering of ErCry4a and reduced motional freedom of ErCry4a associated to the lipid membrane compared to the solution phase.

To further characterize the membrane binding of ErCry4a, the computationally modeled system was employed to analyze the electrostatic potential on the surface of the protein and the membrane patch. The ErCry4a/bilayer system we modelled is illustrated in Fig. 4A. The system was studied using Molecular Dynamics simulations (MD). After extensive equilibration, the average area per lipid was 0.55 nm^2^, and the membrane thickness was ∼4 nm, see Fig. S8. These parameters are characteristic for a liquid disorder state of the lipid membrane.^104^ Figure 4B shows that ErCry4a has a dipole-like distribution of surface charges with a positive pool located closer to the C-terminus (blue color in Fig. 4B) and the negative pool at the N-terminus (red color in Fig. 4B). Due to the two distinctly different charge distributions, two initial orientations of the protein in regard to the membrane were probed to calculate the interaction energy between the protein and the membrane. Figure 4C shows that in the case of the Ori1 simulations, with the ErCry4a oriented with its positively charged C terminus toward the membrane surface and in the presence of Mg^2+^ [Ori1+Mg(1) and Ori1+Mg(2) simulations], a strong attraction between the protein and the membrane was observed (the interaction energy becomes profoundly negative), see supporting video S1. The strong interaction of the protein and the membrane is supported by the analysis of the average distance between the membrane surface and the protein, which decreases to 0.2 nm after 50 ns of the simulation, see Fig. S9. In the second scenario, the negatively charged N-terminal side of ErCry4a was placed above the membrane surface (Ori2+Mg simulation). Such orientation of the protein yielded significantly weaker interaction energy (see Fig. 4D and supporting video S2), indicating an unfavorable binding orientation of ErCry4a. Furthermore, the influence of Mg^2+^ ions on ErCry4a binding to the membrane was probed. The red line in Fig. 4E shows that two different replica simulations yielded different results when Mg^2+^ ions were absent. Even in the favorable Ori1 orientation, in one of the simulations, the interaction energy between the protein and the membrane increased over the simulation time, indicating that ErCry4a failed to bind reliably to the membrane between the protein and the membrane. In that case, the protein started to reorient itself above the membrane surface but made no contact with the lipids (see supporting video S3). However, in another independent simulation performed at identical conditions, see the blue line in Fig. 4E, the binding of ErCry4a to the membrane was possible. Experimental results demonstrated that in the absence of Mg^2+^ ions, ErCry4a binds to the membrane surface, see Fig. 2D. The amount of the adsorbed protein was larger than in the absence of Mg^2+^, and independent experiments exhibited some divergency of the measured absorbance of the amide I’ mode. These results indicate that the Mg^2+^ ions may play some unspecific role in the lipid-protein interaction. To elucidate the role of Mg^2+^ ions in the ErCry4a binding to the membrane, the interaction energy of each Mg^2+^ ion with the membrane and with the protein was computed. Figure S10 illustrates that during the simulation, Mg^2+^ ions did not interact simultaneously with the ErCry4a and the membrane, which suggests that Mg^2+^ ions are not necessarily involved in ErCry4a binding to the membrane. The time evolution of the secondary structure elements in ErCry4a during the membrane association process is shown in Fig. S11. A loss of helicity favoring turns and random coils was observed when the negatively charged side of ErCry4a faced the membrane surface, see Fig. S11G. This result suggests that the electrostatic repulsion between ErCry4a and the membrane may be strong enough to disrupt the protein’s native structure. The C-terminal side in ErCry4a has a net positive surface charge and, thereby, may be responsible for the protein association with the membrane. For Ori1, the simulation results showed increased helices content in ErCry4a; see Fig. S11A-D and Fig. S12. On average, the α-helical content increased from 39% to 49% during 200 ns of the MD simulation, which agrees well with the experimentally determined 50 ± 2 % α-helical content for ErCry4a in the solution phase, see Fig. 2A.

**Figure 4.**
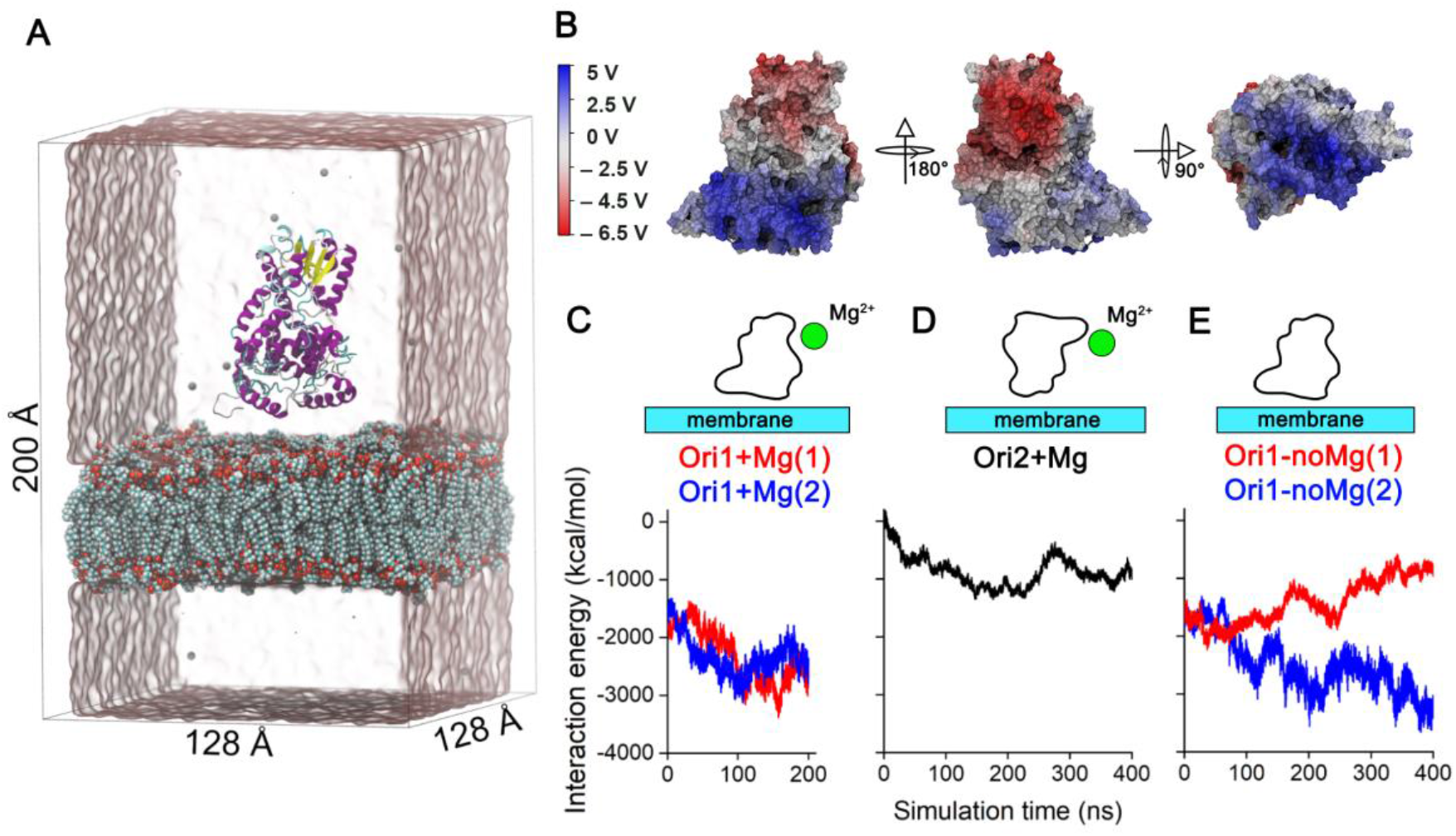
**A**: Depiction of the simulated ErCry4/lipid bilayer system displayed without Na^+^ and Cl^−^ ions for clarity. Magnesium ions (Mg^2+^) are shown as spheres, ErCry4a protein is shown in cartoon representation. Water is illustrated as a transparent surface and the model membrane is shown through van der Waals spheres. **B**: Electrostatic potential of ErCry4a mapped to the surface of the protein. The accompanying scales display the potential in volts. The potential was averaged over the whole production trajectory. **C:** Time evolution of the interaction energy between the ErCry4a protein and the model membrane in three different simulations; **D:** Ori2+Mg **E**: Ori1-noMg scenarios (See Table 1).

To further compare the experimental and computation results, spatial analyses of the α- helices location in the membrane-associated ErCry4a was performed. Specifically, the order parameter of the α-helical fragments of ErCry4a in the membrane-associated state was computed. First, the orientation of the C=O bonds in the helical fragments of the protein with respect to the membrane’s surface normal was determined; the transition dipole moment of the ϖ(C=O) mode is directed along the C=O bond. For each carbonyl group, the angle *θ* between the transition dipole vector and the membrane normal, see Fig. 4A, was established. Averaging *θ* values over all carbonyl groups and over the simulation time results in the average value 〈𝜃〉 that described the coupling of the C=O vibrations to the electric field vector. The computed 〈𝜃〉 value of 43° was used to calculate *S*0 from Eq. (3), which gave a value of 0.60 ± 0.05. This value agrees well with the experimentally determined order parameter of 0.47 ± 0.07 in the presence of Mg^2+^, see Fig. 3. For 0.45<*S*0< 0.60, one expects a specific orientation of the protein in the membrane bound state with a certain degree of motional freedom.

The MD simulations revealed changes in the α-helical content of the C-terminal tail of ErCry4a upon its association with the membrane. The random coil part of the C-terminal tail underwent a conformational change to an α-helix that itself inserted into the polar head group region of the membrane’s outer leaflet, as shown in Fig. S13. In this regard, specific residues in ErCry4a are important, pointing to the formation of hydrogen bonds between the protein and the lipids in the membrane. Figure 5 depicts key amino acid residues engaged in hydrogen bonding (Arg415, Arg497, Lys472, Lys509, and Arg510), supporting the finding that the ErCry4a binding is driven predominantly by electrostatic interactions. Residues 490-527 constitute the C-terminal tail of ErCry4a.^39, 35^ These 37 amino acid residues include 4 arginines, 4 lysines, 5 aspartic acids, and 4 glutamic acids, meaning that 45% of all residues in the C-terminal are charged. This suggests that the C-terminal could be an important protein part that interacts with other cellular components like lipids. Lipids involved directly in the interaction with specific amino acid residues in ErCry4a were identified, see Fig. S14. DMPC, DMPE, and DMPS made contacts predominantly to the positively charged amino acids in the C-terminal tail (residues 490-527). Considering the low content of DMPS (χ= 0.15) in the membrane, this lipid exhibits the most frequent contacts to the C-terminal of ErCry4a, see Table S3 and Fig. S14. Most hydrogen bonding interactions occurred between the positively charged amino acids, mainly Lys and Arg, and the negatively charged carboxylate moiety in DMPS.

**Figure 5:**
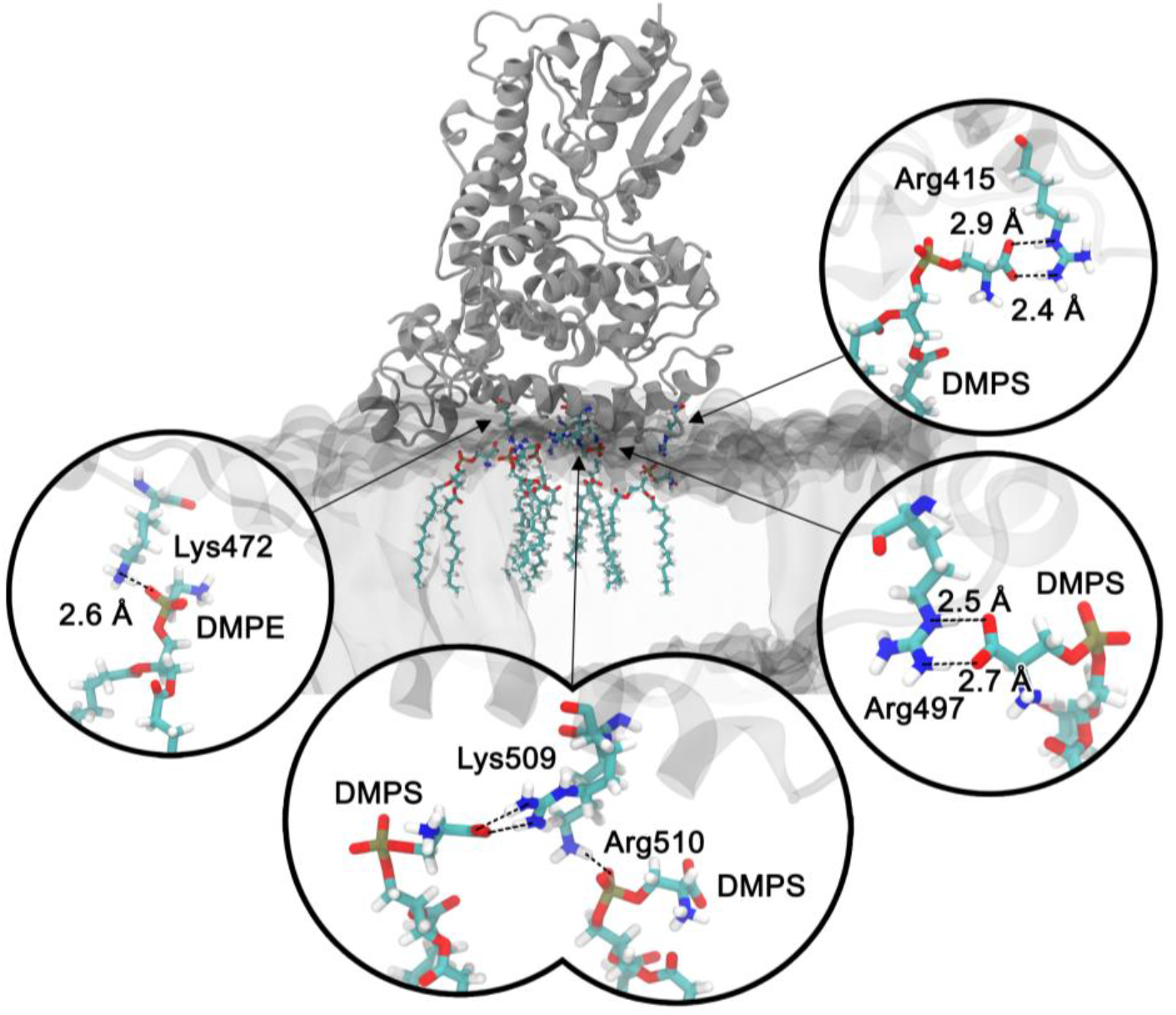
Close up of interactions between the two phospholipids’ (DMPS and DMPE) charged polar heads and charged amino acids located at or close to the C-terminal tail of the ErCry4a protein, highlighting the identified hydrogen bonds.

## Conclusion

Our results showed unambiguously that ErCry4a binds to model membranes simulating outer segments of vertebrate photoreceptor cells. Figure 6 illustrates the invagination of ErCry4a into the membrane, as seen in the MD simulations. The lipid-ErCry4a interaction is dominated by electrostatics and yields uniformly oriented protein molecules on the membrane surface. The results of the atomistic MD simulations indicate that positively charged amino acids at the C terminus tail make hydrogen bonds with PS and PC lipids, embedding the protein into the polar head group region of the membrane. A comparative study that examined the C-terminal regions of plant and animal Cry proteins revealed high flexibility and a propensity to a disordered structure.^105^ The study further showed a light dependent conformational change of the C-terminus in *Arabidopsis* Cry1, which seems to be a hallmark in other Cry forms as well, for example in pigeon ClCry4^47^, chicken ^106^ and drosophila^107^. Wu *et al.* reported that the 60 amino acid long C-terminal fragment of ErCry4a has the potency to interact with the three transmembrane proteins identified as binding partners: long-wavelength sensitive opsin (iodopsin), voltage-gated potassium channel subunit Kv8.2 and retinal G protein coupled receptor. However, the three cytoplasmic putative interaction partners of ErCry4a: guanine nucleotide-binding protein (Gt) subunit alpha.2, guanine nucleotide-binding protein subunit gamma 10 and retinol binding protein 1 did not interact with the isolated C-terminus.^49^ This observation agrees with the notion of Görtemaker *et al.*^51^ that one of the cytoplasmic proteins, the α-subunit of the heterotrimeric cone-specific G protein, can interact with ErCry4a truncated just before the C-terminus. These earlier results indicate different interaction sites in ErCry4a for different interacting proteins and might point to conformational re-arrangements of the C-terminus. The results of the present work also hint at a switch mechanism involving the C-terminus, which is either attached to a lipid membrane and/or engaged in interacting with the retinal transmembrane proteins identified by Wu *et al.*, (e.g., long wavelength sensitive opsin the K^+^-channel subunit Kv8.2, and the retinal G protein-coupled receptor, RGR).^40^ Change of the conformation of a polypeptide chain in a protein when making contact with the lipids is a characteristic feature of the lipid-protein interactions.^108^ This is the first study to show that membrane association could be one of the biological functions of the C-terminal tail in ErCry4a. Although membrane association is thus far not typically considered a primary biological function of the C-terminal tail in other cryptochromes, it is worth noting that some studies have suggested a potential association of cryptochromes with cellular membranes under certain conditions.^109^ For example, in Drosophila, the C-terminal tail of cryptochrome has been implicated in its interaction with a membrane protein called NinaC. However, the precise functions of the C-terminal tail of cryptochromes varies depending on the specific organism and cryptochrome isoform.^105–107^ Membrane association may be a specific function of Cry4.

**Figure 6:**
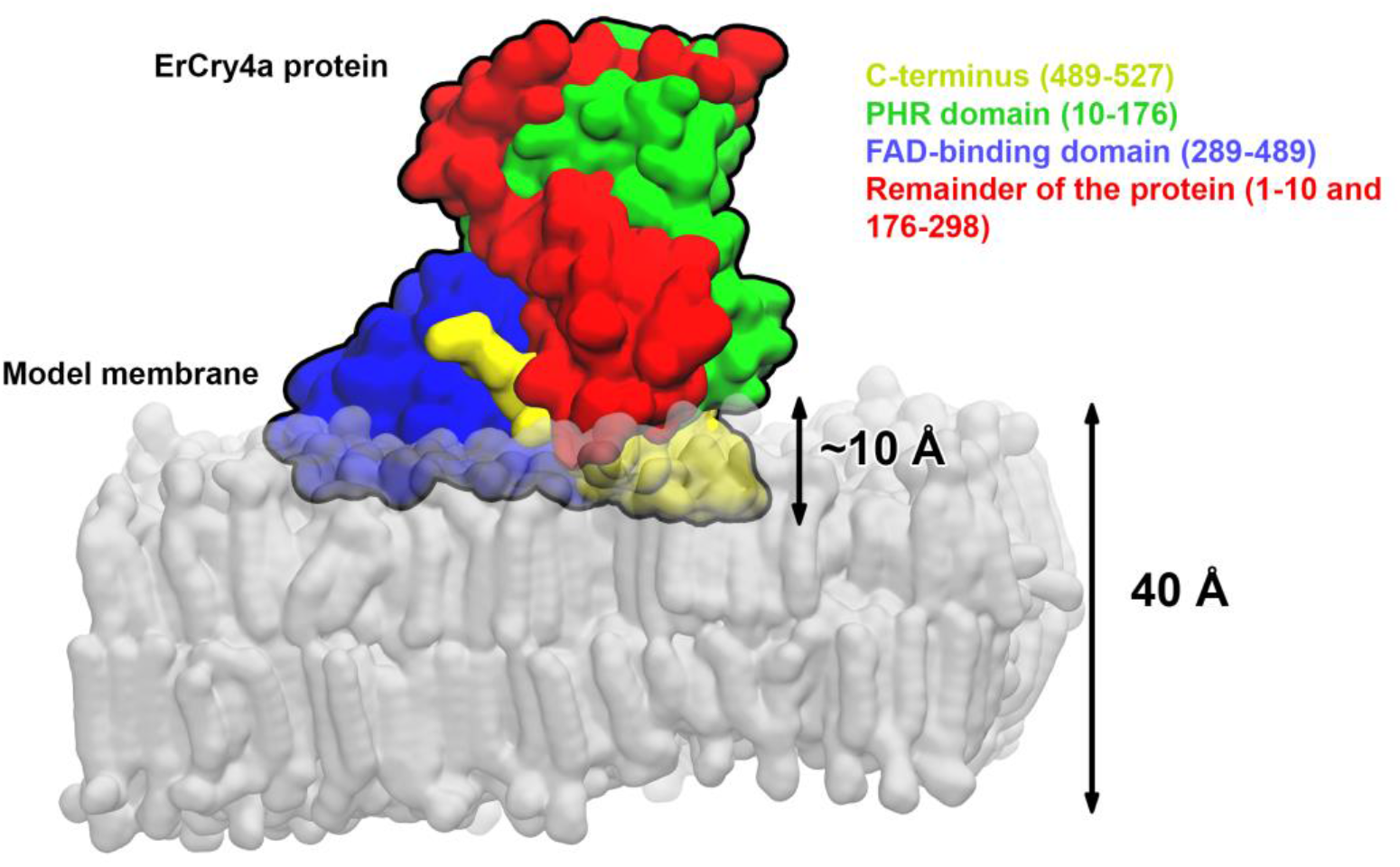
The location and orientation of ErCry4a bound to the model membrane of outer segments of the double cone photoreceptor cells. Different fragments of the protein are marked in the corresponding colors.

**Figure 7:**
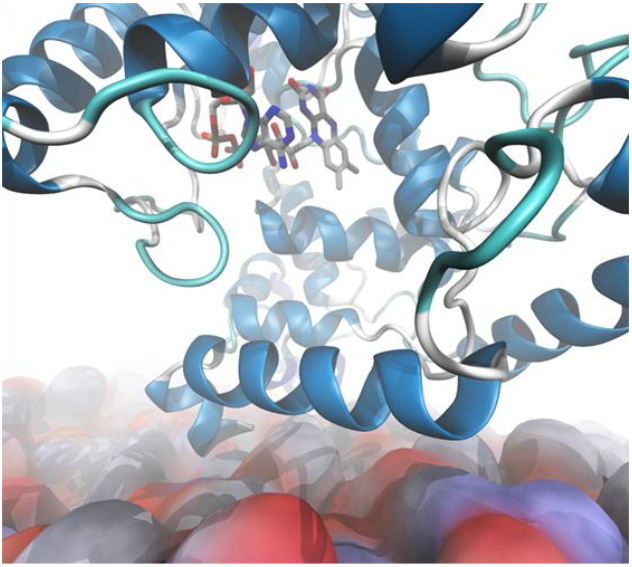
For Table of Contents Only

The quantitative analysis of the amide I’ mode of the experimental PM IRRA spectra revealed the order parameter of 0.47 for the α-helices in the membrane-bound ErCry4a. At the same time, a value of 0.60 was established computationally. Both values are significantly higher than 0.0, expected for a random protein orientation on the membrane surface. However, the found order parameter differs from 1 and −0.5, which is expected for an immobilized protein on the membrane surface. It is, therefore, natural to conclude that ErCry4a has a well-defined orientation in its membrane-associated state, and the motion of the protein is partially restricted. The insertion of ErCry4a into the membrane, see Fig. 6, and its directional binding via the C-terminal reduce the translational mobility of the protein. The protein, however, may still rotate, rock, and wobble at the surface. Orientational order and alignment of helical fragments in ErCry4a implement ordering and reduced mobility of the FAD moiety bound to the protein, and thus, ordering and reduced mobility of the radical pair formed after absorption of blue light by the protein. A moderate reduction of the order parameter has theoretically been predicted for a radical pair buried in a protein which exhibit a response to the Earth’s magnetic field at physiological temperatures.^21, 31^ It was proposed explicitly that *“an order parameter of approximately 0.5 is therefore not a tight constrain on the ordering of the magnetoreceptor molecules”*.^31^ The order parameter of the α-helical fragments in ErCry4a measured and modelled here matches the theoretical requirements for efficient sensing of weak magnetic fields predicted in ref. 31. The results of this combined experimental-theoretical study provide the first experimental indication that ErCry4a protein has the necessary degree of ordering upon interaction with the membrane surface to detect weak magnetic fields.

In the absence of Mg^2+^, the experimentally determined order parameter of α-helices was 0.23 ± 0.07, indicating an increased molecular-scale disorder and a larger mobility of the membrane-associated ErCry4a protein. All experiments featured more intense amide I’ modes once Mg^2+^ was added. Furthermore, the amide I’ modes contained a contribution from aggregated β-sheets, suggesting the accumulation of large amounts of the protein on the membrane surface.^100^ In the absence of Mg2+, two independent simulations yielded different results showing both ability and disability of ErCry4a to bind to the lipid membrane. It is, however, interesting that on one simulation, the orientation of ErCry4a did not differ between the Mg^2+^-free and Mg^2+^-bound states. Combining the experimental and computational results, we speculate that Mg^2+^ ions screen the negative charge at the N-terminus of ErCry4a, preventing the adsorption of multilayers and protein misfolding (denaturation).

At the level of the membrane-protein supramolecular assembly, the MD simulations suggest that the protein protrudes into the membrane’s outer leaflet by ca. 1 nm, see Fig.6. Such a deep insertion of the protein into the polar head group region of the lipid leaflet requires a certain flexibility and elasticity of the membrane. The MD results suggest that ErCry4a binds exclusively to membranes in a liquid disorder state. In this phase, the acyl chains have a significant fraction of gauche conformations, which introduce disorder in packing and give mobility to the lipid molecules.^94^ In the liquid state, the membrane is thinner than a solid state (acyl chains have fully stretched all-trans conformation). A fully stretched myristoyl chain, present in lipids used in this study, is 1.79 nm long.^110^ Adding 0.8 nm for the thickness of the polar head group^111, 112^, a solid-state model lipid bilayer would be ∼5.20 nm thick. The simulated membrane has a thickness of 4.00 nm and corresponds to membranes existing in a liquid-disordered state.^104^ A loose packing of the acyl chains ensures water accumulation in the polar head group region which affects the membrane elasticity.^59^ In summary, the results suggest that the membrane’s physical state governs a cytoplasmic protein’s ability to associate and bind to the membrane surface. Temperature is the driving force for phase transitions in lipid assemblies. Therefore, subtle changes in the thermotropic membrane properties may control the *in vivo* association of ErCry4a to the model of the outer segment double cone photoreceptor membranes.

In conclusion, the membrane lamellae in the outer segments of double-cone photoreceptor cells may serve as an attachment platform for ErCry4a. Cryptochrome ordering on the membrane surface is, however, not the only necessary condition for efficient magnetoreception. In other words, a radical pair and thus a protein have to be aligned within the array of the receptor cells.^21, 31, 35^ It was recently found that double cone photoreceptor cells in European robin display order and regularity.^35^ It is therefore confirmed at the cellular^35^ and molecular level that the ordering degree required for sensing anisotropic Earth’s magnetic fields could be achieved. Verification of ordering of ErCry4a to the outer segment membranes *in vivo* is still required. A further critical aspect of ErCry4a association with membranes is the longitudinal and rotational diffusion on the membrane surface. Future work needs to address whether ErCry4a is assembled with protein complexes, leading to a restriction of diffusion and a more stable anisotropy.

## Supporting information

SupportingFile

## Supporting Information document available

The supporting information contains additional data and information to support our results and interpretations. The document includes the following sections: S1. Composition of the model membrane of the outer segment cone of the vertebrate photoreceptor cell; S2. Preparation of the floating membrane on the gold surface; S3. Equilibrium spreading pressure of ErCry4a; S4. Langmuir isotherms for the lipid mixtures; S5. Secondary structure elements of ErCry4a in solution and in the membrane-associated state; S6. Quantitative analysis of the amide I’ mode of membrane-associated ErCry4a; S7. Correlation between the intensity of an IR absorption band in IRRAS and the orientation of a molecule in an anisotropic film; S8. Physical characteristics of the simulated model membrane; S9. Primary structure analysis of the ErCry4a protein; S10. Distance analysis between the ErCry4a protein and the model membrane; S11. Interaction energy analysis between Mg^2+^ ions, the model membrane, and the ErCry4a protein; S12. Timeline analysis of ErCry4a protein secondary structure changes; S13. Analysis of ErCry4a secondary structure changes; S14. Secondary structure changes of the C-terminal tail in ErCry4a upon interaction with the model membrane; S15. Hydrogen bonds between the model membrane and ErCry4a, and S.16. Hydrogen bond analysis in the MD simulations. Supporting videos S1-S3 illustrate the dynamical behavior of ErCry4a in the vicinity of the membrane surface for the Ori1+Mg (supporting video S1), Ori2+Mg (supporting video S2) and Ori1-noMg (supporting video S3) simulations.

## Acknowledgments

The authors would like to thank the Hanse Institute for Advanced Study (post-doctoral fellowship granted to MM), the Volkswagen Foundation (Lichtenberg professorship awarded to IAS), the Deutsche Forschungsgemeinschaft (GRK1885 Molecular Basis of Sensory Biology to HM, IAS, KWK; SFB 1372 Magnetoreception and Navigation in Vertebrates, no. 395940726 to HM, IAS, KWK; Hyp*Mol-Hyperpolarization in molecular systems, no. 514664767, TRR386/1-2023 to IAS), the Ministry for Science and Culture of Lower Saxony (Simulations Meet Experiments on the Nanoscale: Opening up the Quantum World to Artificial Intelligence, SMART; and Dynamik auf der Nanoskala: Von kohärenten Elementarprozessen zur Funktionalität, DyNano to IAS), and the European Research Council (under the European Union’s Horizon 2020 research and innovation programme, grant agreement no. 810002, Synergy Grant: ‘QuantumBirds’, awarded to HM). Computational resources for the simulations were provided by the CARL Cluster at the Carl-von-Ossietzky University, Oldenburg, supported by the DFG and the Ministry for Science and Culture of Lower Saxony. The authors also gratefully acknowledge the computing time granted by the Resource Allocation Board and provided on the supercomputer Lise and Emmy at NHR@ZIB and NHR@Göttingen as part of the NHR infrastructure. The calculations for this research were conducted with computing resources under the project nip00058.

## Author Contributions

The research idea originated from collaborative work between MM, IB, IAS, HM and KWK. RB and JS produced and purified the protein. MM performed biophysical experiments, including fabrication of the floating lipid membrane, cryptochrome binding to model membrane, IR spectroscopy studies, analyses of the raw data, and wrote the corresponding experimental parts of the manuscript, MH performed all of the computational simulation and analysis, IB performed quantitative analysis of the amide I mode; HM, KWK, MH, IAS and IB wrote and corrected the manuscript.

